# Lung endothelial niche signaling governs self-renewal and fate transitions of human alveolar stem cells

**DOI:** 10.64898/2026.03.07.710316

**Authors:** Bang-Jin Kim, Dasom Hwang, Jimyung Park, Sung Joo Jang, Jaewon Kim, Chiara Camillo, Elena Floris, Austin Choi, Soomin Ryu, Frank D’Ovidio, Sandra Ryeom

**Affiliations:** Division of Surgical Science, Department of Surgery, Columbia University Irving Medical Center, New York, NY, 10032, USA; Department of Medicine, Columbia University Irving Medical Center, New York, NY, 10032, USA; Lung Transplant Program, Columbia University Irving Medical Center, New York, NY, 10032, USA

## Abstract

Chronic lung diseases such as pulmonary fibrosis are characterized by the irreversible loss of alveolar type 1 (AT1) cells, yet the mechanisms governing human alveolar stem cell self-renewal and differentiation remain poorly defined. Here, we identify a lung endothelial niche that sustains the self-renewal of human alveolar type 2 (AT2) stem cells through MAPK signaling, enabling robust long-term expansion while preserving stem cell fate. Although YAP activation initiates AT1 transcriptional programs, it is insufficient to complete lineage maturation. We show that MAPK inhibition together with LATS inhibition promotes nuclear translocation of YAP, enhancing AT1 differentiation. Expanded human AT2 stem cells engraft in fibrotic lungs and contribute to alveolar regeneration while undergoing directed differentiation within diseased human lung tissue. Together, our findings define a niche-controlled signaling mechanism governing human alveolar stem cell fate and advance our understanding of alveolar regeneration.

## Introduction

The lung exhibits a remarkable capacity for regeneration after injury, as demonstrated in cases such as influenza and COVID-19. This regenerative ability is primarily attributed to alveolar type 2 (AT2) cells, which both self-renew and differentiate into alveolar type 1 (AT1) cells^1, 2^. While AT1 cells are essential for gas exchange, AT2 cells serve as a stem cell population in the lung, making them crucial for maintaining alveolar integrity and repair. In pulmonary fibrosis, the damage is typically irreversible with anti-fibrotic drugs only able to slow disease progression leaving lung transplantation as the only treatment option for advanced disease. A promising alternative approach involves the expansion and transplantation of AT2 cells into fibrotic lungs to restore damaged AT1 populations. This has been a successful approach in mouse models of pulmonary fibrosis however, studies of long-term expansion of human AT2 cells have been limited^3^. The growth factors and signaling pathways that maintain human AT2 cell identity and direct differentiation into AT1 cells remain poorly defined.

Here, we identify lung endothelial niche signals that govern human AT2 stem cell self-renewal and fate decisions. Although several studies have reported conditions that promote the self-renewal of human AT2 cells, maintaining their purity and identity over extended periods remains challenging. Adult stem cells are profoundly influenced by their microenvironment with niche - derived signals regulating both self-renewal and differentiation. Lung endothelial cells (LECs) are recognized as active regulators of epithelial repair, modulating injury responses and shaping stem-cell behavior. Mouse studies show that endothelial cues can limit fibrosis, direct bronchioalveolar stem cells (BASCs) toward alveolar lineages, and interact with perivascular fibroblasts that exacerbate fibrotic scarring^4, 5^. These findings underscore the importance of endothelial niche signals in coordinating lung injury and repair. However, the full repertoire of endothelial-derived factors that govern human lung homeostasis and AT2 cell-fate decisions are not fully understood.

Basal cells, which reside in the conducting airways and give rise to secretory and other luminal cell lineages, also impact disease progression^6^. In pulmonary fibrosis, both the extent of fibrotic scarring and the expansion of metaplastic basal cells within the alveolar air sacs, which obstruct gas exchange, are associated with poor patient outcomes^7^. Recent work has shown that metaplastic basal cells can arise from AT2 cells in alveolar air sacs in mouse and 3D organoid models, with fibroblasts providing cues that further promote this aberrant differentiation^8^. Thus, therapeutic strategies must inhibit AT2-to-basal metaplasia while directing AT2 cells towards AT1 differentiation.

Yes-associated protein (YAP) is a transcriptional co-activator and the principal effector of the Hippo pathway, modulating gene expression in response to mechanical and biochemical signals to govern organ size and tissue homeostasis^9, 10, 11^. Upon nuclear accumulation, YAP activates lineage-specific transcriptional programs, making it a pivotal regulator of stem- and progenitor-cell fate decisions during differentiation^12, 13, 14^. Recent studies highlight a key role for YAP in alveolar epithelial fate: in mouse models, YAP loss increases AT2 marker expression, whereas YAP activation enhances chromatin accessibility at lineage-defining loci and induces AGER, a hallmark AT1 gene^15^. In another study, human AT1 transcriptome analysis identified Hippo–LATS–YAP/TAZ signaling as a defining feature of AT1 cells, with LATS1/2 inhibition driving iPSC-derived AT2 cell differentiation into AT1-like cells^16^. These data suggest that YAP is a central regulator of AT1 differentiation across species. Despite the clear importance of YAP in promoting AT1 differentiation, it remains uncertain whether YAP-driven programs can generate the level of AT1 regeneration required for functional repair in the human air sac.

In this study, we show that human lung endothelial signals acting through MAPK sustain human AT2 self-renewal and that inhibiting this pathway enables their directed differentiation into AT1 cells, addressing a central barrier in human lung regeneration.

## Results

### A lung endothelial niche governs human AT2 stem cell self-renewal

To define niche signals governing human AT2 stem cell self-renewal, we first established a high-purity AT2 isolation strategy. We utilized a previously reported method based on differential adhesion to collagen I to enhance the purity of human AT2 cells^17^. Collagen-based enrichment followed by HT2-280 sorting yielded highly pure human AT2 populations, as confirmed by robust Surfactant Protein-C (SP-C) expression (Fig. 1a–b). To identify niche cells supporting AT2 self-renewal, we co-cultured HT2-280+ AT2 cells with adult lung fibroblasts (ALFs) and LECs isolated from healthy human lungs. ALF and LEC identities were confirmed by immunostaining for vimentin and CD31, respectively (Supplementary Fig. 1). Co-culture experiments showed that LECs significantly enhanced AT2 expansion, yielding the highest number of SP-C–positive cells, whereas ALF co-cultures increased basal cell differentiation as indicated by Keratin 5 (KRT5) expression. In contrast, the AT1 differentiation marker AGER did not show significant differences among the groups (Fig. 1c–d). These data are consistent with previous studies indicating that adult fibroblasts can promote AT2 differentiation towards KRT5+ basal cells^8^. To understand the paracrine signals responsible, we analyzed ALF and LEC conditioned media using a growth factor array, revealing significantly higher secretion of EGF, PDGF-AA, and TGF-α by LECs (Fig. 1e, Supplementary Fig. 2). We next evaluated these candidate growth factors alongside additional factors known to be crucial in organoid and stem cell cultures, including Noggin, R-spondin, Wnt3a, and FGF10. Our data demonstrate that EGF, FGF10, and TGF-α individually most effectively promoted AT2 proliferation (Fig. 1f). Among various combinations of these growth factors, the EGF + FGF10 combination was most effective in driving AT2 proliferation (Fig. 1g). Western blot analysis confirmed downstream signaling activation, evidenced by increased phosphorylation of ERK and AKT following treatment with these factors alone or in combination with LEC-conditioned media (Supplementary Fig. 3). Pharmacological inhibition of PI3K and MAPK pathways markedly suppressed AT2 cell proliferation (Supplementary Fig. 4a) and analysis of AT2 cultures revealed that the addition of either inhibitor significantly reduced SP-C-expressing AT2 cells. Notably, MAPK inhibition led to a significant increase in double positive SP-C⁺/AGER⁺ cells, indicating a shift toward AT1 differentiation, whereas PI3K inhibition resulted in a significant increase in KRT5⁺ cells, indicating basal cell differentiation (Supplementary Fig. 4b–d). Together, these findings define distinct roles for MAPK and PI3K signaling in human AT2 fate decisions, with MAPK inhibition promoting AT1 differentiation and PI3K inhibition enabling differentiation toward a KRT5-positive basal cell fate. To evaluate whether these optimized growth factor conditions could sustain AT2 identity in three-dimensional culture, we embedded AT2 cells within Matrigel domes using conventional organoid methods^18^. Expansion of KRT5+ basal cells in the culture reduced AT2 culture purity and longevity (data not shown), identifying differentiation towards a KRT5^+^ basal lineage as a barrier to long-term AT2 maintenance. We therefore examined whether intrinsic cellular properties could distinguish basal-prone populations from self-renewing AT2 cells. Notably, KRT5+ basal cells together with KRT5+/SP-C-low and KRT5-/SP-C-double-negative populations exhibited minimal adhesion to Matrigel-coated surfaces, whereas SP-C-positive AT2 cells preferentially bound to Matrigel (Fig. 2a–b). Leveraging this difference, we implemented a Matrigel-based adhesion strategy to selectively retain AT2 cells while eliminating non-AT2 populations. To determine whether selective depletion of basal-prone cells altered regenerative capacity *in vivo*, we transplanted Matrigel-adherent and non-adherent cell populations into bleomycin-injured mice. Matrigel-adherent AT2 cells formed markedly fewer and smaller basal cell colonies whereas non-adherent cells generated extensive KRT5-expressing basal colonies following transplantation (Fig. 2c). Together, these findings show that selective removal of basal-like cells by selecting Matrigel-adherent cells supports AT2 maintenance, thus this selection step was applied at every passage for all subsequent culture steps.

**Fig. 1:**
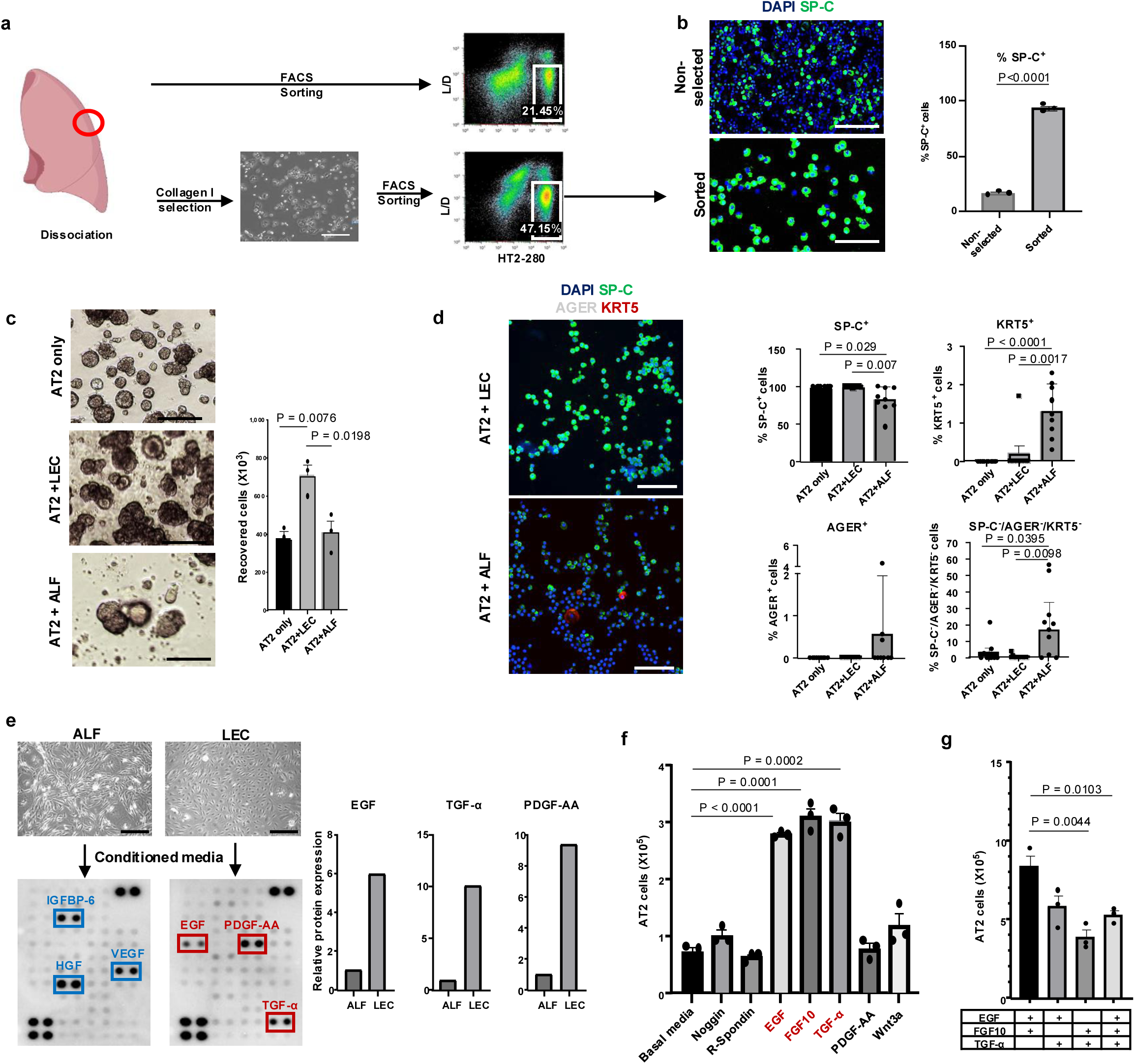
Lung endothelial niche signals enable long-term expansion of human AT2 stem cells. a. Representative images showing the selection process to enrich AT2 cell purity. Scale bar = 300 µm. b. Immunofluorescence staining of SP-C in unsorted lung cells and cells sorted using collagen I and HT2-280 antibody. Scale bar = 100 µm. The graph shows the percentage of SP-C⁺ cells (n = 3). c. Phase-contrast images after co-culture of indicated cells with AT2 cells. Scale bar = 200 µm. The graph shows the number of recovered cells (n = 3). d. Immunofluorescence images of SP-C, AGER and KRT5 in co-cultured cells and recovered cells. The graph shows the percentage of cells expressing each marker protein (n = 9). e. Phase images of adult lung fibroblast (ALF) and lung endothelial cell (LEC) and growth factor array blots using ALF- or LEC-conditioned medium. Scale bar = 250 µm. Quantification of indicated growth factors from ALF and LECs are shown on the right. Protein expression levels are presented as relative values, normalized to the ALF group. f–g. AT2 cell counts after culture with the indicated growth factors (n = 3). Statistical significance was determined by Student’s t test, and data are represented as mean ± SEM.

**Fig. 2:**
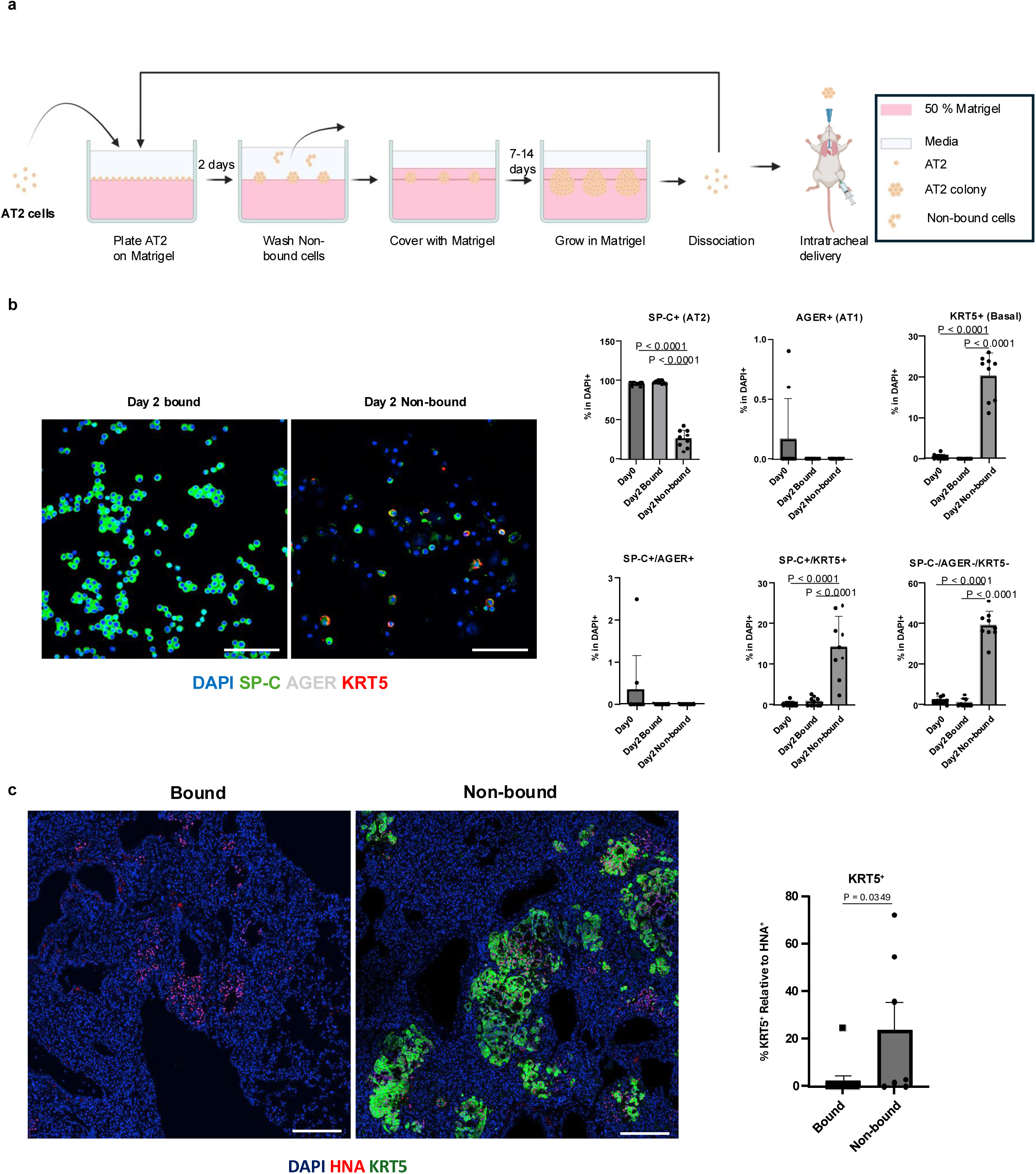
Matrigel-based selection enriches human AT2 stem cells and suppresses KRT5⁺ basal differentiation in vivo. a. Schematic of the feeder-free culture system. b. Immunofluorescence staining of SP-C, AGER, and KRT5 in Matrigel-bound versus non-bound cells. Scale bar = 100 µm. Graphs show the percentage of cells expressing each marker protein (n = 9). c. Immunofluorescence images of human neutrophil antigen (HNA) and KRT5 staining of lungs after transplantation of Matrigel-adherent and non-adherent cells. Graphs show the relative frequency of KRT5^+^ cells relative to HNA expression. Scale bar = 200 µm. Non-selected (n = 7), Selected (n = 11). Statistical significance was determined by Student’s t test, and data are represented as mean ± SEM.

### Long-term cultured human AT2 Cells maintain stable progenitor identity

We evaluated maintenance of AT2 cell identity during long-term expansion by analyzing cells at multiple passages (P0, P5, P12). Anti-SP-C immunofluorescence demonstrated sustained, high-level SP-C expression across all passages with minimal expression of AGER and KRT5 (Fig. 3a). Single-cell RNA sequencing analyses further validated these findings by mapping our cultured cells onto UMAP plots derived from published datasets of healthy human lungs, specifically epithelial cell populations. Cell-annotation algorithms identified 95% AT2 cells at P0, 99% at P5, and 91% at P12 in our cultured cells (Fig. 3b). Additionally, gene expression profiling showed robust and consistent expression of AT2-specific markers *SFTPC* and *surfactant protein B (SFTPB)*, with minimal expression of differentiation markers *AGER* and *KRT5* (Fig. 3c–d). Collectively, these comprehensive analyses confirm the effectiveness of our optimized culture strategy in maintaining AT2 cells and minimizing differentiation towards a basal cell fate.

**Fig. 3:**
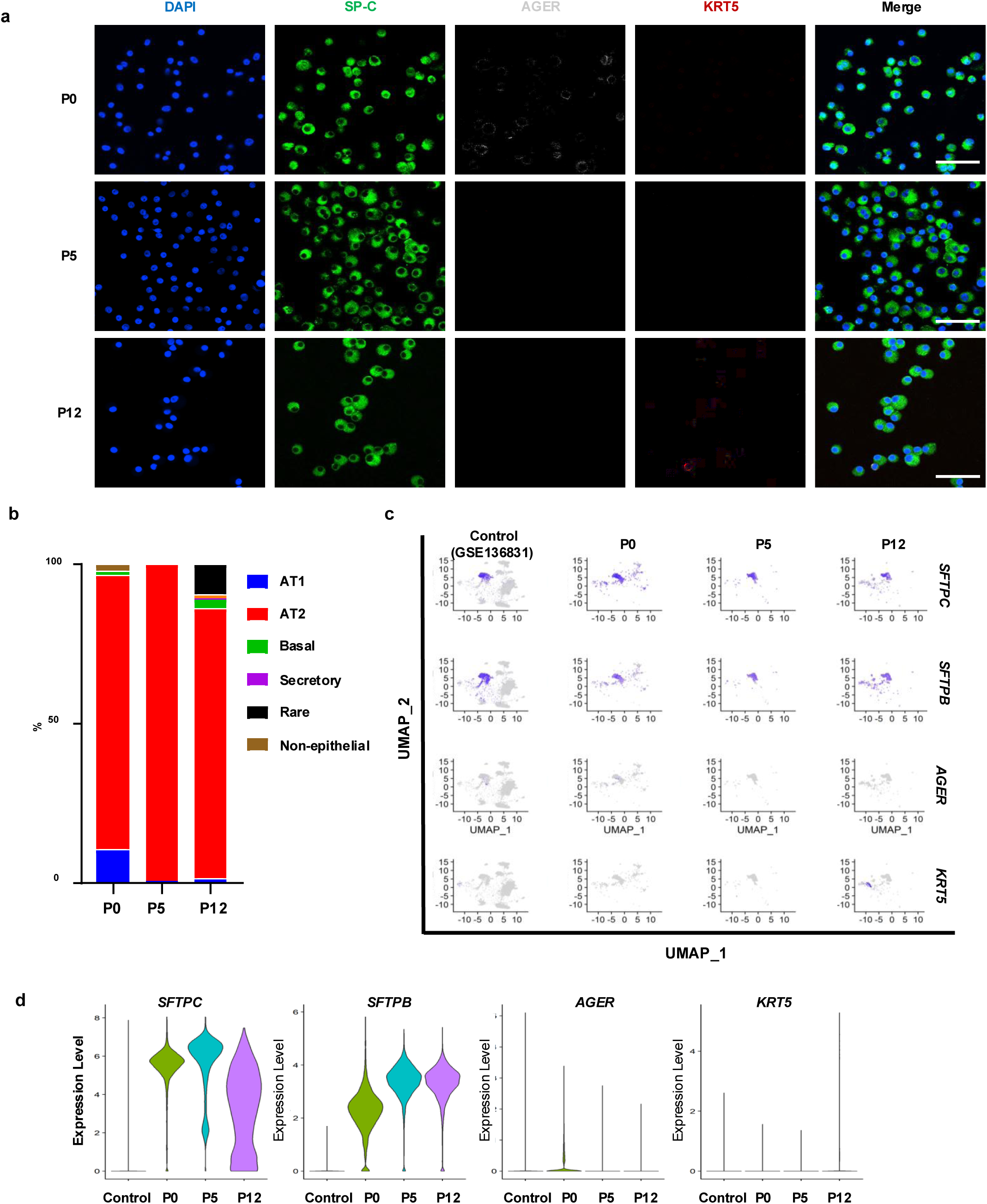
Long-term cultured human AT2 cells maintain their identity. a. Immunofluorescence of SP-C, AGER, and KRT5 in long-term cultured AT2 cells at the indicated passages. Cells were dissociated into single-cell suspensions, cytospun onto slides, and processed for immunofluorescence analysis. (P). Scale bar = 50 µm. b–d. Single-cell RNA sequencing analysis of AT2 cells across passages. b. Cell-type annotation by passage. c. Expression patterns of AT2 markers (*SFTPC*, *SFTPB*, *AGER* and *KRT5*) in UMAP plots across passages. d. Violin plots showing expression of *SFTPC*, *SFTPB*, *AGER* and *KRT5*.

### YAP activation initiates the AT1 program

While prior studies, largely in mouse and iPSC-derived systems, have shown that promoting YAP nuclear localization drives AT2-to-AT1 differentiation, this has not been directly examined in primary human AT2 cells. Thus, we treated human AT2 cells with a large tumor suppressor kinase (LATS) inhibitor for one week to assess whether inhibition of LATS signaling promotes differentiation toward an AT1 cell fate. LATS inhibitor–treated colonies displayed significant expansion in size (Fig. 4a). Western blot analyses confirmed reduced YAP phosphorylation, decreased SP-C expression, and increased AGER expression in LATS inhibitor-treated cells (Fig. 4b). Immunofluorescence analysis confirmed that LATS inhibition promoted YAP nuclear translocation, with a significant increase in YAP nuclear localization compared to controls ( Fig. 4c). Further, qPCR revealed increased expression of the YAP target ankyrin repeat domain 1 (*ANKRD1*), reduced expression of *SFTPC* and lysosome-associated membrane glycoprotein 3 (*LAMP3*), and elevated expression of AT1-specific markers including *AGER*, Rhotekin 2 (*RTKN2*) and caveolin-1 (*CAV1*) (Fig. 4d). Nearly half of the LATS-treated cells expressed both SP-C and AGER (Fig. 4e). This observation suggests that although LATS inhibition effectively initiates the differentiation program, it primarily drives cells into an intermediate AT1 state expressing both AT1 and AT2 markers rather than fully mature AGER-positive, SP-C-negative AT1 cells.

**Fig. 4:**
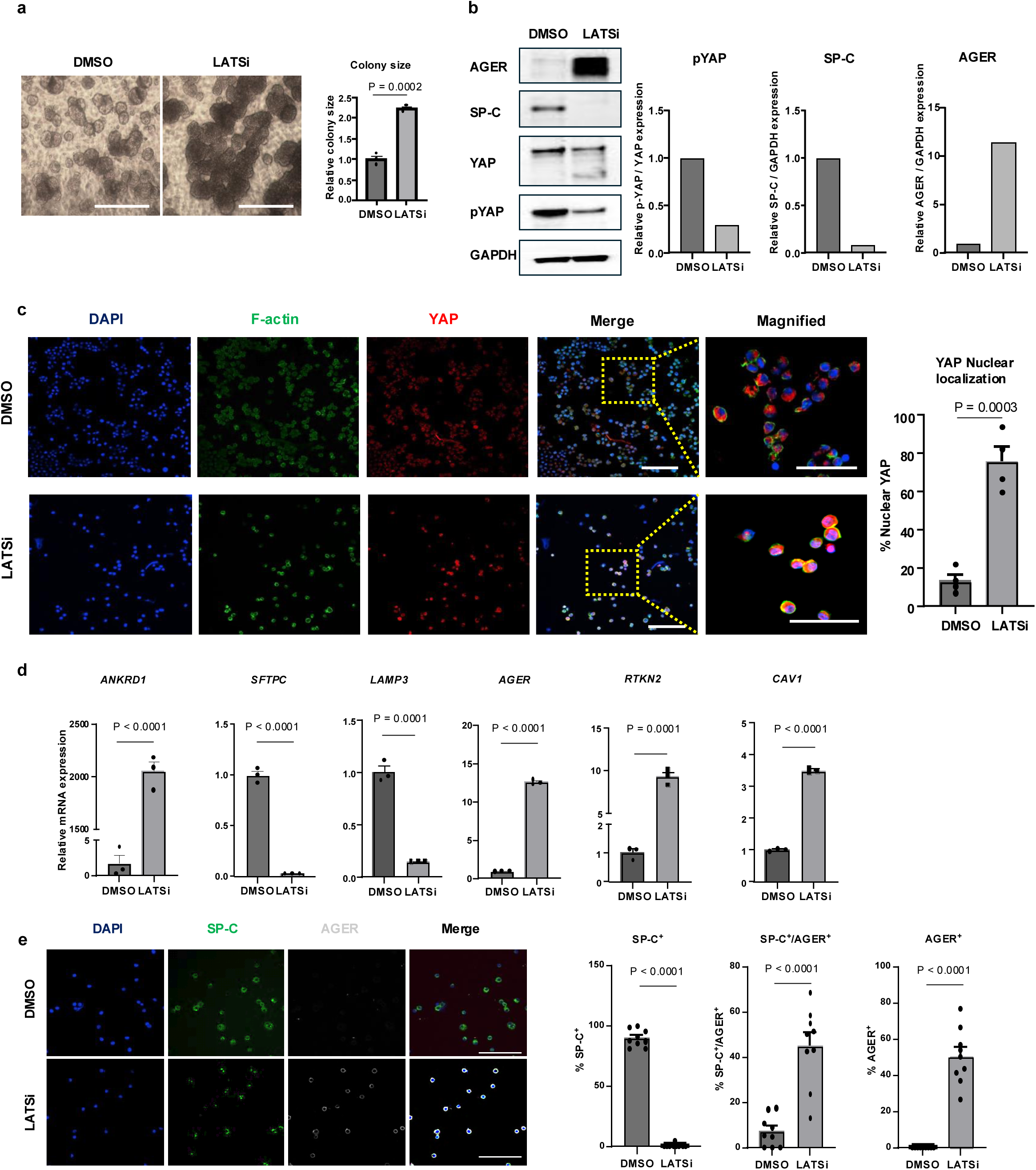
LATS inhibition promotes differentiation toward an AT1 fate. a. Phase images of cells cultured for 1 week. Scale bar = 300 µm. The graph shows relative colony size (n = 3). b. Western blot analysis of AGER, SP-C, YAP, and phosphorylated YAP (pYAP) in cells after the indicated treatment. Graphs indicate relative protein expression levels. c. Immunofluorescence staining of F-actin and YAP in cells following the indicated treatment. Cells were dissociated into single-cell suspensions, cytospun onto slides, and processed for immunofluorescence analysis. Scale bar in merged images = 100 µm, magnified = 50 µm. The graph shows the percentage of YAP nuclear localization (n = 4). d. Relative mRNA expression of *ANKRD1*, *SFTPC*, *LAMP3, AGER*, *RTKN2* and *CAV1* in recovered cells (n = 3). e. Immunofluorescence of SP-C and AGER in cells after indicated treatment. Scale bar = 100 µm. The graph shows the percentage of cells expressing each protein (n = 3). Statistical significance was determined by Student’s t test, and data are represented as mean ± SEM.

### Long-term cultured human AT2 Cells engraft and differentiate in a fibrotic mouse lung

We next examined whether long-term *in vitro* expanded human AT2 cells retain the capacity for AT1 differentiation following transplantation into a fibrotic lung environment. To this end, human AT2 cells were transplanted into immunodeficient mice with bleomycin-induced lung fibrosis to assess their engraftment and differentiation potential *in vivo*. Human nuclear antigen (HNA) staining was used to identify transplanted human cells within the mouse lung tissue. LATS inhibition significantly increased the number of HNA-positive engrafted human cells, expanded the size of human cell–derived colonies, and promoted differentiation toward a transitional AGER⁺ AT1-like cell state *in vivo* (Fig. 5a–b). Notably, LATS inhibitor–treated lungs also contained increased numbers of SP-C⁺ human cells, suggesting that YAP activation supports both AT2 persistence and progression toward the AT1 lineage within the injured lung (Fig. 5c).

**Fig. 5:**
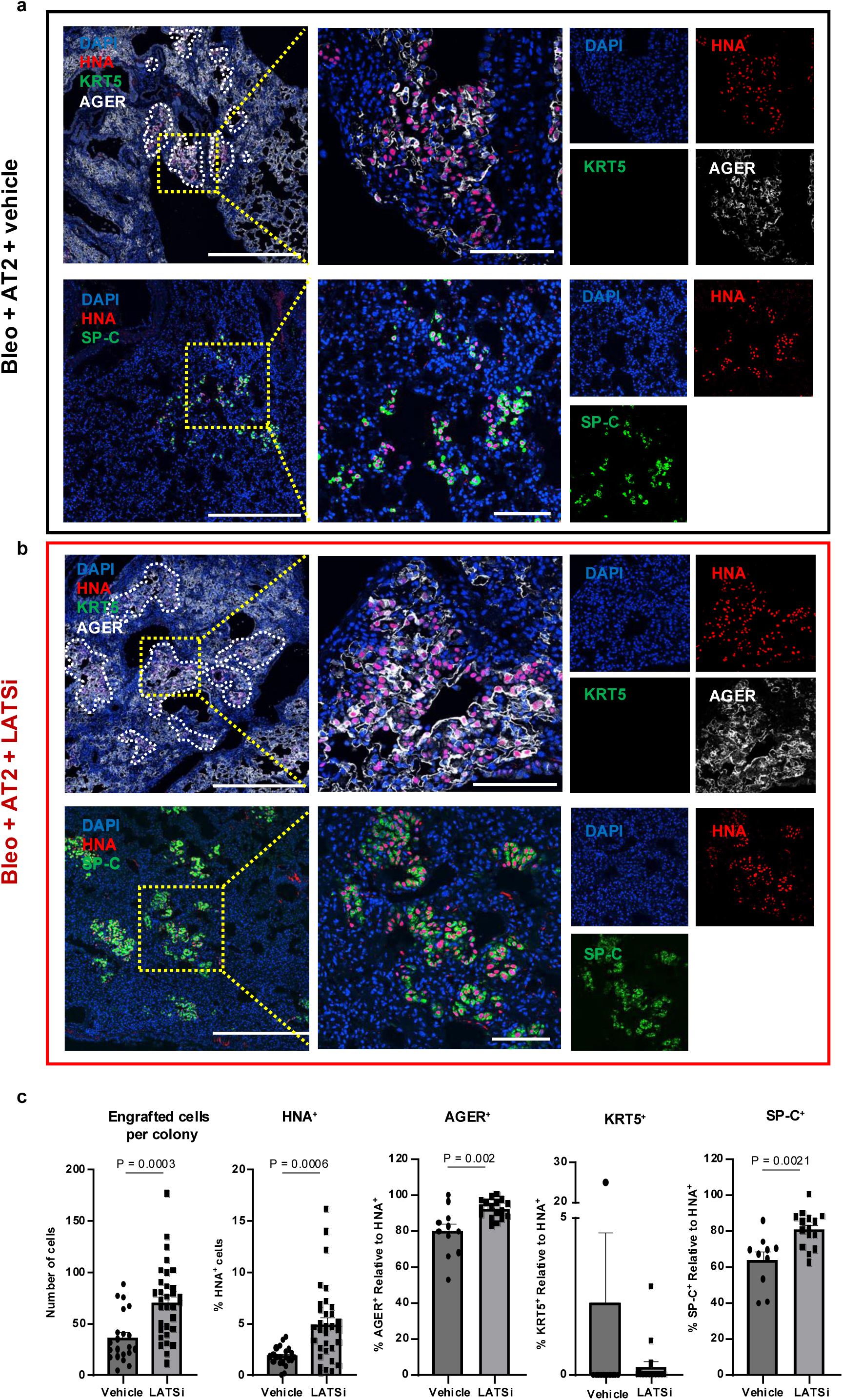
Long-term cultured human AT2 cells engraft and differentiate in a mouse model of fibrosis. a–b. Immunofluorescence of KRT5/AGER (co-staining) and SP-C in lung sections from each mouse group. Human nuclear antigen (HNA) staining was used to detect human cells. a. Lung images from mice inoculated with AT2 cells only. b. Lung images from mice inoculated with AT2 cells and treated with LATS inhibitor. Scale bars: whole-scan images = 500 µm, magnified = 100 µm. c. Graphs show number of engrafted cells per colony after indicated treatment as well as the percentages of HNA⁺ cells and AGER^+^, KRT5^+^ or SFTPC^+^ cells relative to HNA expression. Statistical significance was determined by Student’s t test, and data are represented as mean ± SEM.

### MAPK inhibition and YAP activation enhance AT2-to-AT1 differentiation

Collectively, our data suggest that persistent AT2 self-renewal signaling constrains differentiation toward an AT1 fate. Given that MAPK/ERK activity sustains AT2 stem cell identity and that MAPK inhibition increases the proportion of AGER⁺ cells, we hypothesized that combined inhibition of MAPK and LATS would further promote differentiation toward an AT1 fate. Combined inhibition of LATS and MAPK markedly reduced SP-C⁺ cells and significantly increased AGER⁺ single-positive AT1-like cells as compared to inhibition of either pathway alone (Fig. 6a). MAPK inhibition alone increased the transitional SP-C⁺/AGER⁺ populations, whereas dual inhibition promoted progression to AGER+ and SP-C-negative cell state. qPCR analysis corroborated these findings, revealing the lowest *SFTPC* expression and strongest *AGER* induction under combined pathway suppression (Fig. 6b). Together, these results identify MAPK signaling as a barrier to complete AT1 differentiation and demonstrate that coordinated modulation of MAPK and YAP pathways is required for efficient resolution of the AT1 fate program.

**Fig. 6:**
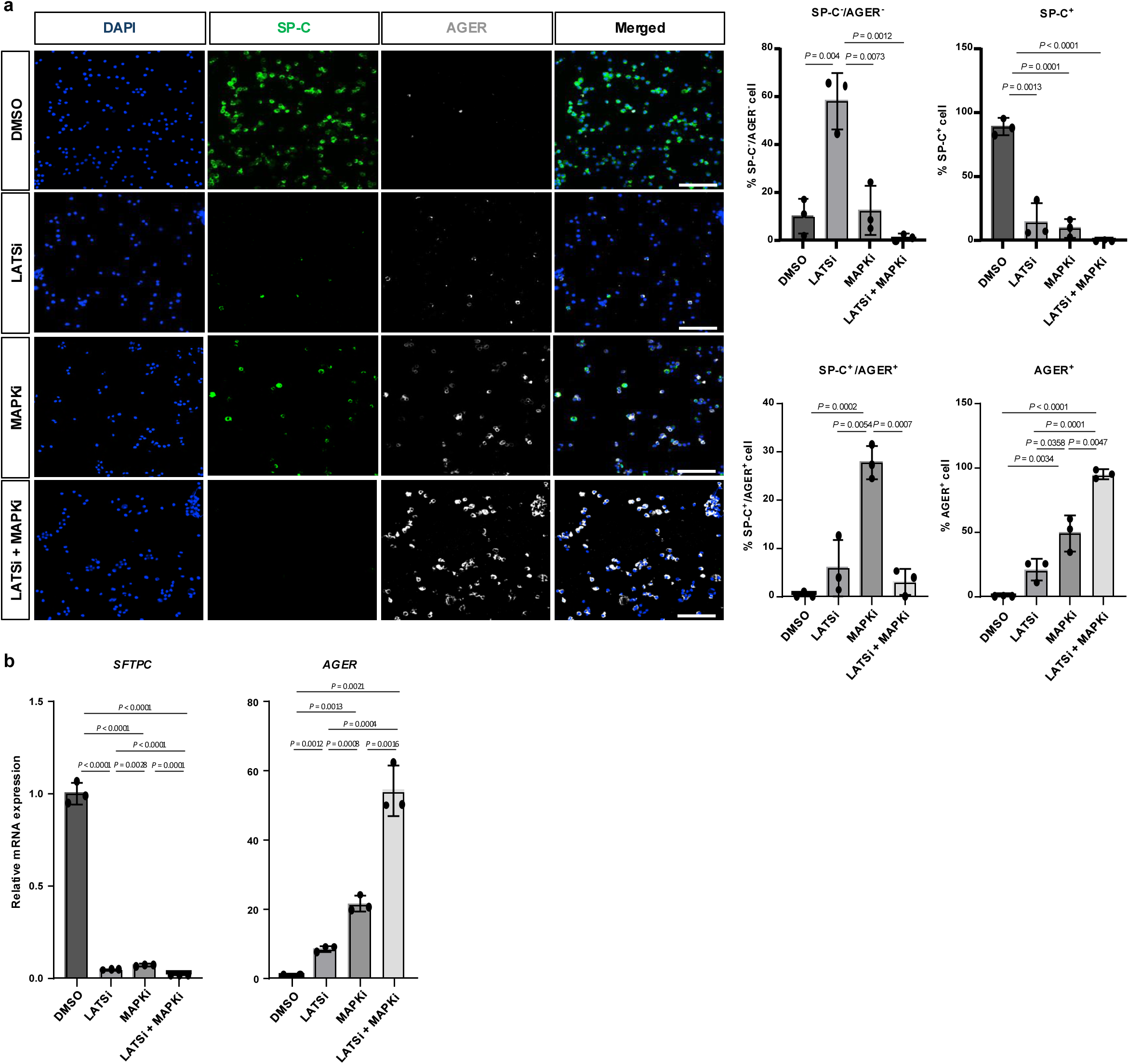
MAPK inhibition and YAP activation drive AT2-to-AT1 differentiation. a. Representative immunofluorescence images of SP-C and AGER in the indicated treatment groups. Bar graphs on the right show the percentage of cells in each population, including SP-C⁻/AGER⁻, SP-C⁺, SP-C⁺/AGER⁺, and AGER⁺. b. Relative mRNA expression levels of SP-C and AGER in cells harvested after 12 days after the indicated treatment. Scale bar = 100 µm. Graph shows the percentage of cells expressing each protein (n = 3). Statistical significance was determined by Student’s t test, and data are represented as mean ± SEM.

### Combined LATS and MAPK inhibition enhances AT2-to-AT1 differentiation in human fibrotic lung slices

To determine whether dual inhibition of MAPK and LATS promotes AT1 differentiation within a human fibrotic microenvironment, we utilized precision-cut lung slices (PCLS) derived from patients with pulmonary fibrosis (Fig. 7a). GFP-labeled AT2 cells cultured with fibrotic PCLS (Fig. 7b) exhibited higher SP-C expression in the DMSO vehicle group compared to drug-treated conditions, indicating maintenance of an AT2 phenotype under basal conditions. In contrast, LATS inhibition increased the proportion of GFP⁺/AGER⁺ cells, with an even greater expansion of GFP⁺/AGER⁺ double-positive cells observed under combined LATS and MAPK suppression (Fig. 7c–d), consistent with enhanced differentiation toward an AT1-like lineage. Notably, KRT5 expression was higher in the DMSO group compared to LATS inhibitor–treated cells and was absent in the combined LATS+MAPK condition, suggesting suppression of basal-like differentiation upon coordinated pathway modulation.

**Fig. 7:**
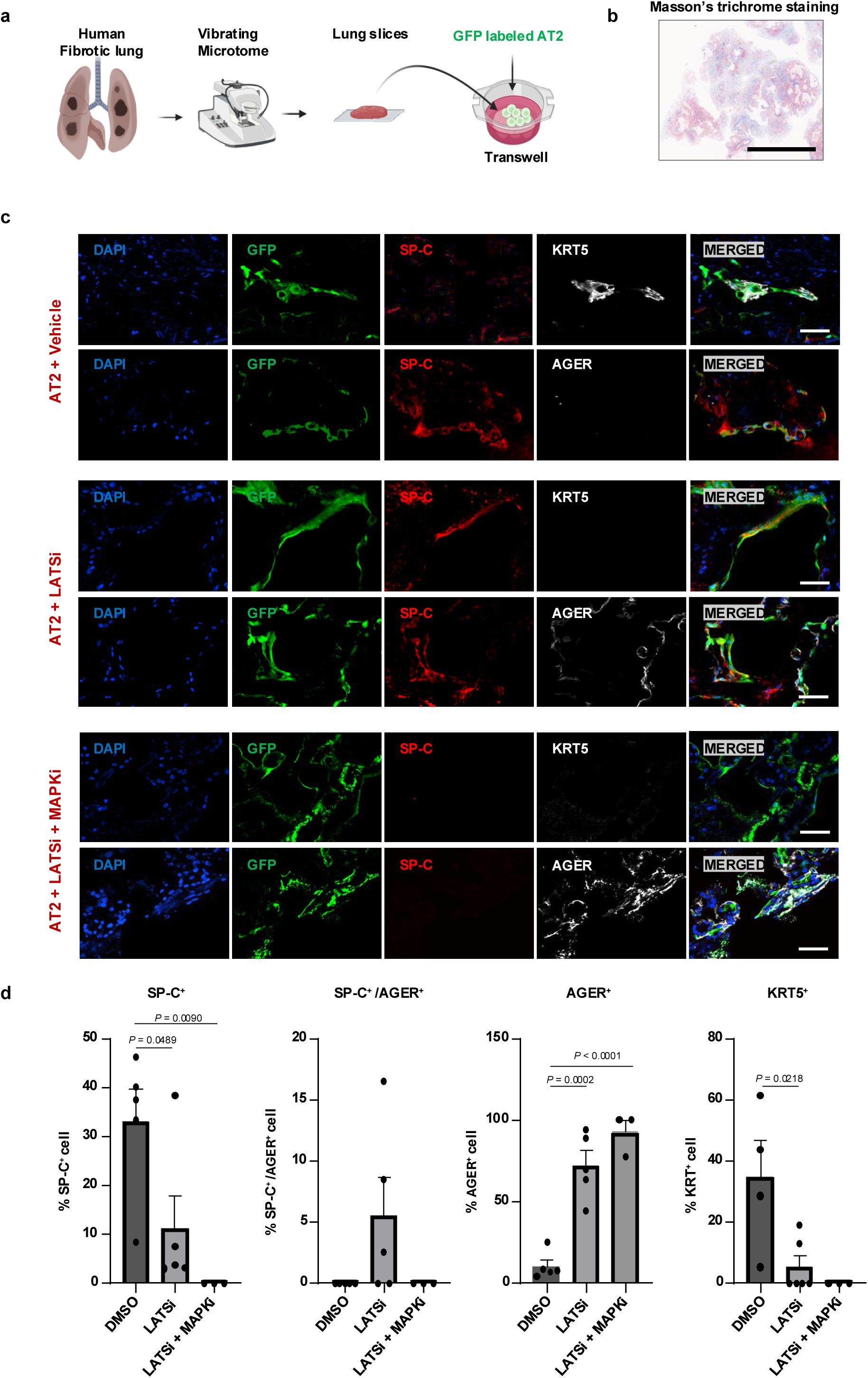
Dual inhibition of LATS and MAPK promotes AT2-to-AT1 differentiation in human fibrotic lung slices. a. Schematic illustration of human AT2 cell transplantation into human fibrotic precision cut lung slices (PCLS). b. Masson’s trichrome staining confirms extensive collagen deposition and fibrotic architecture in human lung tissue used for PCLS experiments. Scale bar = 3 mm c. Immunofluorescence staining of GFP, SP-C, KRT5, and AGER in PCLS following the indicated treatments. For each treatment condition, two representative images are shown: the upper row displays co-staining of GFP, SP-C, and KRT5 to assess basal cell differentiation, whereas the lower row displays co-staining of GFP, SP-C, and AGER to evaluate AT1 differentiation. GFP marks transplanted human cells, and nuclei are counterstained with DAPI. Merged images are shown in the right panels. Scale bar = 50 µm. d. Graphs show the percentage of cells expressing SP-C, KRT5 and AGER in each group. Sample sizes for quantification were as follows: %SP-C⁺: DMSO (n=5), LATSi (n=5), LATSi+MAPKi (n=3); %SP-C⁺/AGER⁺: DMSO (n=5), LATSi (n=5), LATSi+MAPKi (n=3); %AGER⁺: DMSO (n=5), LATSi (n=5), LATSi+MAPKi (n=3); %KRT5⁺: DMSO (n=4), LATSi (n=6), LATSi+MAPKi (n=3). Statistical significance was determined by Student’s t test, and data are represented as mean ± SEM.

## Discussion

Here, we define critical factors derived from a lung endothelial niche and signaling pathways that govern human alveolar stem cell fate, addressing a central barrier to alveolar regeneration in pulmonary fibrosis. Irreversible loss of functional AT1 cells is a defining feature of fibrotic lung disease, yet strategies to sustain human alveolar stem cells while enabling productive differentiation remained limited. By identifying niche-derived signals that preserve AT2 identity and delineating pathways that release constraints on lineage progression, our study identifies key pathways governing human alveolar stem cell fate.

Stem cell behavior is fundamentally shaped by microenvironmental cues that coordinate self-renewal and lineage specification. Our findings position lung endothelial cells as a critical niche population supporting human AT2 proliferation, whereas adult lung fibroblasts bias differentiation toward basal and AT1 states, consistent with prior reports^8^. Our findings indicate that ERK and PI3K signaling serve as central regulators of human AT2 fate, jointly sustaining self-renewal while exerting distinct influences on lineage commitment. Specifically, ERK inhibition promotes AT1 differentiation, whereas PI3K inhibition drives cells toward a basal lineage, underscoring the divergent roles of these pathways in directing AT2 cell fate. These pathways therefore represent key nodes for balancing AT2 maintenance with directed differentiation, providing a conceptual framework for promoting alveolar regeneration in fibrotic lung disease. Although MAPK inhibition significantly reduced AT2 proliferation in our study, MAPK blockade has previously been shown to attenuate pulmonary fibrosis by suppressing fibroblast activation and TGF-β signaling^19^. Notably, these anti-fibrotic studies have relied on receptor-level modulators such asα₂-adrenergic or ADORA2B-acting compounds, whereas our study utilized the MEK inhibitor Trametinib to directly suppress ERK signaling and interrogate epithelial lineage control. Collectively, these mechanistically distinct findings support the rationale for using MEK-targeted MAPK inhibition in our differentiation experiments to promote AT1 differentiation within a fibrotic milieu.

Prior studies implicate high TGF-β–expressing stromal cells in promoting AT2-to-basal cell differentiation, a process associated with poor patient survival^8^. TGF-β mediates its effects via both SMAD-dependent and independent pathways, notably through activation of the PI3K–AKT–mTOR axis^20, 21^. In our study, pharmacological inhibition of PI3K increased the proportion of basal cells. These data underscore the distinct signaling axes governing human AT2 lineage self-renewal versus differentiation. This paradoxical effect suggests that PI3K signaling may, in certain contexts, function as a negative regulator of basal lineage commitment, potentially by providing feedback inhibition to SMAD activation or by restraining alternative pro-differentiation pathways. Further mechanistic studies will be required to elucidate the precise role of PI3K in this process, which may ultimately enable strategies to prevent AT2-to-basal cell conversion in fibrotic lung disease.

Beyond intracellular signaling pathways, we identified a functional property that distinguishes distinct cellular states during long-term culture. Basal-like and SP-C-low cells exhibited minimal adhesion to extracellular matrix substrates, whereas self-renewing AT2 cells preferentially attached to Matrigel. This differential adhesion enabled selective depletion of basal-like populations and improved the stability of the AT2 cultures during expansion. Importantly, failure to remove these cells resulted in extensive basal colony formation following transplantation, whereas Matrigel adherent cell preparations showed enhanced AT2 engraftment and expansion. Together, these findings suggest that functional heterogeneity emerges during prolonged culture and that matrix adhesion-based selection strategies can help maintain alveolar epithelial homeostasis.

Our findings demonstrate that YAP activation and MAPK suppression act through complementary mechanisms to drive AT2 cells toward an AT1 fate. LATS inhibition, which enhances YAP activity, strongly induced AT1-associated transcriptional programs but predominantly generated intermediate or transitional AT1-like cells expressing both AT1 and AT2 markers. These data indicate that YAP activation alone is insufficient to complete the differentiation trajectory towards AT1 cells. In contrast, MAPK inhibition enhanced progression toward an AGER⁺ AT1-like phenotype, yielding significantly higher AGER⁺ populations than LATS inhibition alone and revealing an ERK-dependent barrier that limits terminal AT1 differentiation by human AT2 cells. The markedly enhanced expansion of AT1 cells observed with dual LATS and MAPK inhibition suggests that coordinated modulation of both pathways is required to overcome this barrier to drive cells towards an AT1 identity. Targeting both LATS and MAPK also improved AT2 differentiation towards AT1 cells in fibrotic PCLS, indicating that dual regulation of YAP and MAPK signaling can enhance AT1 generation even within a fibrotic microenvironment. Collectively, our findings support a LEC niche-controlled model of fate progression in human alveolar stem cells. Lung endothelial signals sustain MAPK-dependent AT2 progenitor maintenance, whereas modulation of MAPK and YAP activity enables efficient differentiation towards an AT1 fate while limiting differentiation of basal lineages. By establishing principles that govern expansion, preservation, and directed differentiation of human AT2 cells, this work advances fundamental understanding of alveolar regeneration and provides a conceptual foundation for future regenerative strategies targeting the injured lung.

## Methods

### Human lung tissue dissociation

Fresh or frozen-thawed human lung tissue transported on ice was minced into ∼2 mm fragments using fine scissors. Tissue was digested in 5 mg/mL Collagenase type II (Worthington Biochemical, LS004174) and 500 µg/mL DNase I in a volume equivalent to three times the tissue volume, incubated at 37 °C, 250 rpm for 30–40 min. The digested suspension was diluted with an equal volume of ALF medium and filtered through a 100 µm cell strainer. Red blood cells were lysed by adding ACK lysis buffer (Gibco, A10492-01) for 2 min, followed by quenching with ≥10 volumes of ALF medium and centrifugation. The cell pellet was washed and filtered through a 70 µm cell strainer to obtain a single-cell suspension for downstream applications.

### AT2 cell purification

Single-cell suspensions of human lung were prepared as described above and immediately transferred into ALF medium additionally supplemented with gentamicin (0.2 µg/mL; Gibco, 15750060) and amphotericin B (1.25 µg/mL; Gibco, 15290026). Tissue-culture dishes were pre-coated with collagen I (20 µg/mL; Sigma-Aldrich, C3867) for 1 h at 37 °C, washed once with PBS, and equilibrated with ALF medium. Cells were plated onto collagen-coated dishes and incubated overnight at 37 °C, 5% CO₂. The next day, non-adherent cells and debris were removed with medium exchange followed by 3–5 additional washes with PBS (Gibco, 10010-023). Adherent cells were detached with 0.25% trypsin-EDTA (Gibco, 25200072) for 3 min at 37 °C and neutralized with 3 volumes of ALF medium. Cells were pelleted (500 g, 5 min), resuspended in 500 µL PBS-S (PBS containing 1% FBS), and incubated with 25 µL anti-human HT2-280 mouse IgM (Terracebiotech, TB-27AHT2-280) for 15 min on ice. After addition of 10 mL PBS-S, cells were centrifuged (500 g, 5 min), the supernatant was aspirated, and the pellet was resuspended in 500 µL PBS-S. A secondary antibody (goat anti-mouse IgM Alexa Fluor 488, Invitrogen, A21042; 5 µL) and a fixable Live/Dead viability dye (Invitrogen, L34957; 5 µL) were added and cells were incubated for 15 min on ice. Cells were washed with 10 mL PBS-S, centrifuged (500 g, 5 min), resuspended in 10 mL PBS-S, and filtered through a 40-µm cell strainer (Corning, 352340) into a 15-mL conical tube. Cells were counted, pelleted once more (500 g, 5 min), and resuspended in PBS-S to 2–3 × 10^6^ cells/mL (2 mL PBS-S was pre-loaded into a fresh 15-mL tube for sorting). Fluorescence-activated cell sorting was performed on a Sony MA900 sorter. Live single cells were gated as Live/Dead-negative, and alveolar type 2 (AT2) cells were collected as HT2-280-positive events.

### ALF and EC isolation and culture

Single-cell suspensions of human lung tissue were plated on 0.1% gelatin-coated culture dishes (room temperature, 15 min; Sigma-Aldrich, G1890) and incubated for 1 h at 37 °C in pre-warmed ALF medium supplemented with gentamicin and amphotericin B. Adherent cells were subsequently maintained in ALF medium, and cells with passage number ≤ 8 were used for co-culture experiments. Non-adherent cells were collected after 1 h and re-plated on 0.1% gelatin-coated dishes in endothelial cell growth medium (EGM; Lonza, CC-3162) and cultured overnight. For EC purification, adherent cells were harvested, washed 2–3 times with PBS, and detached by trypsinization (Gibco, 25200072). Cells were pelleted (500 g, 5 min), resuspended in 5 mL PBS-S, and passed through LS magnetic columns (Miltenyi Biotec, 130-042-401) to deplete cells containing lung-derived magnetic particles that can cause false-positive CD31 labeling. Flow-through fractions were collected, centrifuged (500 g, 5 min), and resuspended in 500 µL PBS-S. Cells were incubated with 10 µL human Fc receptor (hFCR) blocking reagent (Miltenyi Biotec, 130-059-901) for 5–10 min on ice, followed by 50 µL anti-human CD31 antibody for 15 min on ice. Samples were washed with 10 mL PBS-S, centrifuged (500 g, 5 min), resuspended in 5 mL PBS-S, and reapplied to LS columns. After three washes with 5 mL PBS-S, CD31⁺ endothelial cells were eluted, centrifuged (500 g, 5 min), and seeded on 0.1% gelatin-coated dishes. Medium was changed after overnight incubation, and cells were expanded by serial passaging. Cells with passage number ≤ 8 were used for downstream co-culture experiments.

### AT2 cell culture and expansion

FACS-purified AT2 cells were embedded in Matrigel-based 3D cultures. Complete medium (CM) was mixed 1:1 with Matrigel (Corning, 354234) and 350 µL of the 50% Matrigel mixture was plated per well of a 24-well plate, evenly spread, and solidified for 30 min at 37 °C. Sorted AT2 cells (3-4 × 10^5^ cells/well) were seeded on top in 700 µL CM. After 24-48 h, the medium containing floating cells and debris was carefully aspirated, and adherent cells were washed twice with pre-warmed PBS. A 150-µL overlay of 50% Matrigel was added and allowed to solidify for 30 min at 37 °C, followed by the addition of 700 µL pre-warmed CM. Cultures were maintained with medium exchange three times per week. The complete medium consisted of Advanced DMEM/F12 (Gibco, 12634010) supplemented with Insulin-Transferrin-Selenium (1×; Gibco, 41400045), HEPES (15 mM; Gibco, 15630080), GlutaMAX™ Supplement (1×; Gibco, 35050061), B-27 Supplement minus vitamin A (1×; Gibco, 12587010), N-2 Supplement (1×; Gibco, 17502048), and Antibiotic-Antimycotic (1×; Gibco, 15240062). The following additives were included: SB431542 (10 µM; Abcam, ab120163), CHIR99021 (3 µM; Tocris, 4423), BIRB796 (1 µM; Tocris, 5989), recombinant human EGF (50 ng/mL; R&D Systems, 236-EG), heparin (5 µg/mL; Sigma-Aldrich, H3149), N-acetyl-L-cysteine (1.25 mM; Sigma-Aldrich, A9165), Y-27632 2HCl (10 µM; Selleck Chemicals, S1049), recombinant human FGF10 (20 ng/mL; BioLegend, 559304), and recombinant human Noggin (100 ng/mL; R&D Systems, 3344-NG). Subculture was performed every 7–10 days. Cultures were incubated with Dispase (4 mg/mL in PBS; Sigma-Aldrich, D4693) freshly prepared and filtered, supplemented with DNase I (500 µg/mL; Worthington, LS002139), at 2 mL per well for 40 min at 37 °C with gentle pipetting, followed by centrifugation (500 g, 5 min). Cell pellets were digested with 0.25% trypsin-EDTA containing DNase I (500 µg/mL; Sigma, DN25) for 5 min at 37 °C with pipetting (10–15 times) before quenching with 3 volumes of ALF medium. After centrifugation, cells were re-plated following the procedure described above. For growth factor screening, AT2 cells were seeded into 48-well plates (1 × 10^5^ cells/well) in 50% Matrigel (175 µL/well) and cultured for 7 days. From the basal medium, which excluded all growth factors present in the conditioned medium (CM), we individually reintroduced each factor—Noggin (R&D Systems, 3344-NG), R-Spondin (R&D Systems, 4645-RS), EGF (R&D Systems, 236-EG), FGF10 (BioLegend, 559304), TGFα (R&D Systems, 239-A), and PDGF-AA (R&D Systems, 221-AA)—each at 50 ng/mL, and performed the assays. Parallel conditions were tested in 24-well plates (3 × 10^5^ cells/well) with growth factor combinations for 1 week, and cell numbers were quantified. EGF (50 ng/mL) and FGF10 (20 ng/mL;) were identified as optimal supplements, and all subsequent experiments were performed using this combination. Recombinant Noggin was included only during early passages (up to P2) and was omitted thereafter. To assess the impact on AT2 cell self-renewal, we added the ERK inhibitor Trametinib (10 nM, MedChemExpress, HY-10999) and the PI3K inhibitor Idelalisib (2 µM, Selleckchem, S2226) to the culture medium. For single-cell RNA-seq and in vivo transplantation studies, cultures were maintained on Matrigel as described, washed to remove floating cells, and adherent cells were harvested by sequential dispase and trypsin digestion for downstream applications.

### Co-culture of AT2 cells with ALF or EC

For co-culture experiments, 3 × 10^5^ AT2 cells were embedded in Matrigel in 24-well plates. On top of the AT2 layer, 1 × 10^5^ ALF or ECS were mixed with 30 μL of 50% Matrigel and overlaid, followed by polymerization. Cultures were subsequently imaged for morphological assessment. For experiments requiring selective recovery of AT2 cells, an indirect co-culture system was employed. AT2 cells (3 × 10^5^) were plated in 24-well plates, and ALF or EC (1 × 10^5^ in 70 μL of 50% Matrigel) were seeded in transwell inserts placed above the AT2 cultures. This setup enabled paracrine interactions without direct contact, allowing recovery and downstream analyses (cell counting and immunofluorescence) of AT2 cells only.

### Growth factor array

ALFs and ECs (2 × 10^5^ cells) were plated in 6-well plates and cultured overnight. The following day, cells were washed three times with PBS and switched to serum-free medium. After 48 h, conditioned medium was collected and analyzed using the Human Growth Factor Antibody Array (Abcam, ab134002) according to the manufacturer’s instructions. Signal intensities were quantified and compared using Fiji software.

### Single-cell RNA sequencing and data preprocessing

To determine the cellular composition of cultured human AT2 cells, single-cell RNA sequencing (scRNA-seq) was performed. Cultured cells were collected and submitted as a cell pellet resuspended in PBS with 10% FBS to the Columbia Genome Center, where library preparation and sequencing were performed using the 10x Genomics Chromium Single Cell 3’ v3 Reagent Kit, following the manufacturer’s protocol. Sequencing was conducted on an Illumina NovaSeq 6000 platform. Raw sequencing data were processed and aligned to the human genome (GRCh38) using Cell Ranger (v7.2.0, 10x Genomics) to generate gene-by-cell count matrices for downstream analysis.

### Cell type annotation

To enable comparative analysis, a publicly available human lung scRNA-seq dataset (GSE136831) was downloaded. Only control group samples were extracted, and gene features were matched across datasets using Ensembl gene IDs. The combined dataset was normalized and integrated using the Seurat R package (v5.1.0), followed by dimensionality reduction and clustering. Cell type annotation was performed using the Azimuth platform (https://azimuth.hubmapconsortium.org/) with the Human Lung Cell Atlas (HLCA), which includes 584,944 cells from 107 individuals. The integrated Seurat object was projected onto the HLCA reference UMAP and mapped to annotated lung cell types using Azimuth’s supervised prediction pipeline.

### Code availability

All analyses were conducted using publicly available software and packages, as described in the Methods. No custom code was developed. R scripts used for data analysis and figure generation are available from the corresponding author upon reasonable request.

### Immunofluorescence (IF)

For cytospin preparations, harvested cells were fixed overnight at 4 °C in 2% paraformaldehyde (Electron Microscopy Sciences, 15710), resuspended at 1 × 10^5^cells/100 μL in 70% ethanol, and centrifuged onto Superfrost Plus slides (Fisher Scientific, 12-550-15) using a Shandon Cytospin 4 (Thermo Scientific) at 1,000 rpm for 5 min. Slides were air-dried completely prior to IF. For mouse lung tissue, lungs were carefully excised to preserve the trachea, inflated with 4% paraformaldehyde, and fixed overnight. Samples were embedded in paraffin and sectioned at 3 μm thickness. Paraffin sections were processed by heat incubation (65 °C, 15 min), followed by xylene (3 min) and graded ethanol (100%, 90%, 70%, and 50%). Sections were rinsed in deionized (DI) water, then treated with antigen unmasking solution (2 mL in 200 mL DI water; Vector Laboratories, H3300250) in a pressure cooker (30 min). After cooling to room temperature (10 min) and rinsing under running tap water (5 min), sections were incubated in PBST for 30 min at room temperature and blocked with CAS-Block™ (Thermo Fisher, 008120) for 10 min. Slides were circled using an ImmunoEdge pen (Vector Laboratories, H-4000) and incubated with primary antibodies overnight at 4 °C. After washing with PBST (2 × 10 min), secondary antibodies were applied, followed by PBST washes (2 × 10 min). Nuclei were counterstained and mounted with Fluoro-Gel II with DAPI (Electron Microscopy Sciences, 1798550). Cytospin-prepared slides were processed identically, beginning from PBST incubation.

### Animals

For lung cell transplantation, NOD-SCID male mice (8–12 weeks old) were anesthetized with 3% isoflurane and placed on an intubation stand (Kent Scientific, ETI-MSE-01). Mice received an intratracheal injection of bleomycin (1 IU/kg; MFR Pharmaceuticals, 71288-0101-10) using a mouse intubation kit (BMR Supply, Ohan-201). Ten days later, 1 × 10^6^ cells suspended in 30 μL PBS were delivered intratracheally. Lungs were harvested one-month post-transplantation for immunofluorescence analysis. All animal experiments were performed in accordance with Columbia University Institutional Animal Care and Use Committee (IACUC) guidelines.

### AT2 cell differentiation

To induce AT2 cell differentiation, 24-well Transwell inserts (Corning, 353097) were coated with 70 μL of 50% Matrigel and allowed to solidify for 30 min. A total of 1 × 10^5^ cells were seeded in complete medium onto the inserts, and 700 μL of complete medium supplemented with either DMSO, 10 μM LATS inhibitor (MedChemExpress, HY-138489), 10 nM Trametinib, or a combination of LATS inhibitor and Trametinib was added to the lower chamber. Cultures were incubated overnight, and colony formation was confirmed the following day. The transwell medium was then replaced with 30 μL of 50% Matrigel to overlay the colonies, and cells were maintained for 7-12 days to promote differentiation, with medium changes performed three times per week.

### Human lung slice preparation and culture

A lung lobe was obtained via lobectomy. Human lung tissue was placed on ice immediately after collection and transported to the laboratory. A 2% agarose solution (Sigma-Aldrich, A9045) was injected through the bronchioles using a syringe to inflate the lung, after which the tissue was immersed in PBS at 4 °C for 1 h to allow the agarose to solidify. Cylindrical cores were then generated using a biopsy punch, and precision-cut lung slices (PCLS; 500 μm thickness) were prepared using a vibrating microtome (Compresstome VF-310-0Z). Slices were transferred to 6-well plates, washed, and incubated overnight at 37 °C in PCLS medium (DMEM supplemented with 10% FBS, 1% glutamine, 2% penicillin–streptomycin, 0.2 µg/mL gentamycin, and 1.25 µg/mL amphotericin B). The following day, slices were transferred to 24-well transwell inserts. For cell labeling, GFP lentiviral particles (Santa Cruz, sc-108084) were applied at a multiplicity of infection (MOI) of 5 and incubated overnight. GFP-positive cells were subsequently isolated by FACS and expanded in culture. Expanded cells (1 × 10^5^) were mixed with 50% Matrigel and overlaid onto pre-cooled (4 °C, 10 min) lung slices. The plates were further incubated at 4 °C for 20 min to promote cell sedimentation into the slices, followed by transfer to a 37 °C incubator. PCLS medium containing either DMSO or LATS inhibitor (MedChemExpress, HY-138489) was added to the lower chamber, and slices were cultured prior to downstream analyses.

### Antibodies

For immunofluorescence (IF), the following primary antibodies were used: surfactant protein C/SP-C (H-8) (Santa Cruz Biotechnology, sc-518029; 1:50), RAGE/AGER (R&D Systems, AF1145; 1:100), Cytokeratin 5 [EPR1600Y] (Abcam, ab75869; 1:100), Cytokeratin 5 [XM26] (Abcam, ab17130; 1:100), NuMA (Abcam, ab86129; 1:100), YAP (G-6) (Santa Cruz Biotechnology, sc-376830; 1:100), CD31 (Agilent Dako, M0823; 1:100), and Vimentin (D21H3) XP® (Cell Signaling Technology, 5741; 1:100). F-actin was visualized using Phalloidin–Atto 565 (Sigma-Aldrich, 94072; 1:500). Secondary antibodies included donkey anti-rabbit IgG Alexa Fluor 594 (Invitrogen, A-21207; 1:500), donkey anti-goat IgG Alexa Fluor Plus 488 (Invitrogen, A32814TR; 1:500), donkey anti-mouse IgG Alexa Fluor 647 (Invitrogen, A-31571; 1:500), and donkey anti-rabbit IgG Alexa Fluor 647 (Invitrogen, A-31573; 1:500). For flow cytometry (FACS), cells were labeled with HT2-280 (Terrace Biotech, TB-27AHT2-280; 1:20) followed by goat anti-mouse IgM Alexa Fluor 488 (Invitrogen, A-21042; 1:100). For magnetic-activated cell sorting (MACS), FcR Blocking Reagent, human (Miltenyi Biotec, 130-059-901; 1:50) and CD31 MicroBeads, mouse (Miltenyi Biotec, 130-091-935; 1:10) were used. For western blotting (WB), the following primary antibodies were used: phospho-p44/42 MAPK (Erk1/2) (Thr202/Tyr204) (Cell Signaling Technology, 9101; 1:1000), p44/42 MAPK (Erk1/2) (137F5) (Cell Signaling Technology, 4695; 1:1000), phospho-Akt (Ser473) (Cell Signaling Technology, 9271; 1:1000), and Akt1/2/3 (H-136) (Santa Cruz Biotechnology, sc-8312; 1:1000).

### Quantitative PCR

Quantitative PCR was performed using gene-specific primers as follows: SFTPC (forward 5′-CACCTGAAACGCCTTCTTATCG-3′, reverse 5′-TTTCTGGCTCATGTGGAGACC-3′), AGER (forward 5′-ACTACCGAGTCCGTGTCTACC-3′, reverse 5′-GGAACACCAGCCGTGAGTT-3′), KRT5 (forward 5′-CCAAGGTTGATGCACTGATGG-3′, reverse 5′-TGTCAGAGACATGCGTCTGC-3′), ANKRD1 (forward 5′-CGTGGAGGAAACCTGGATGTT-3′, reverse 5′-GTGCTGAGCAACTTATCTCGG-3′), LAMP3 (forward 5′-ACTACCCCAGCGACTACAAAA-3′, reverse 5′-CTAGGGCCGACTGTAACTTCA-3′), RTKN2 (forward 5′-AGCTCACACTACCCTAACCTTG-3′, reverse 5′-CATGTTGCCATACAGGGGAAG-3′), CAV1 (forward 5′-GCGACCCTAAACACCTCAAC-3′, reverse 5′-ATGCCGTCAAAACTGTGTGTC-3′), and GAPDH (forward 5′-AGCCACATCGCTCAGACAC-3′, reverse 5′-GCCCAATACGACCAAATCC-3′).

### Statistics

All statistical analyses were performed using unpaired two-tailed Student’s t-tests. Graphs and statistical evaluations were generated using Prism 10 (GraphPad Software, San Diego, CA). P values indicating significant differences between groups are shown in the figures.

## Data Availability

The data that support the findings of this study are available from the corresponding author upon reasonable request.

## Acknowledgments

We thank the Penn-CHOP Lung Biology Institute for providing human lung tissues used in this study. We acknowledge the Columbia Stem Cell Initiative Flow Cytometry Core for assistance with cell sorting and analysis and the Sulzberger Genome Center for generating the single-cell RNA sequencing data. This work utilized the Confocal and Specialized Microscopy Shared Resource of the Herbert Irving Comprehensive Cancer Center at Columbia University, supported in part by the NIH/NCI Cancer Center Support Grant P30CA013696.

## Author contributions

B.-J.K. conceived the study, developed the methodology, performed experiments, analyzed the data, and contributed to writing, review and editing of the manuscript.

D.H. performed experiments, analyzed the data, review and editing of the manuscript.

J.P. analyzed the data and editing of the manuscript.

S.J.J., J.K., C.C., E.F., and A.C. performed experiments, review and editing of the manuscript.

F.D. provided resources, review and editing of the manuscript, and supervised the study.

S.R. conceived the study, developed the methodology, analyzed the data, contributed to writing, review and editing of the manuscript, and supervised the study.

## Materials & Correspondence

Correspondence and requests for materials should be addressed to Sandra Ryeom.

