## Supplemental figures for "Lung endothelial niche signaling governs self-renewal and fate transitions of human alveolar stem cells"

**Supplemental Fig. 1**

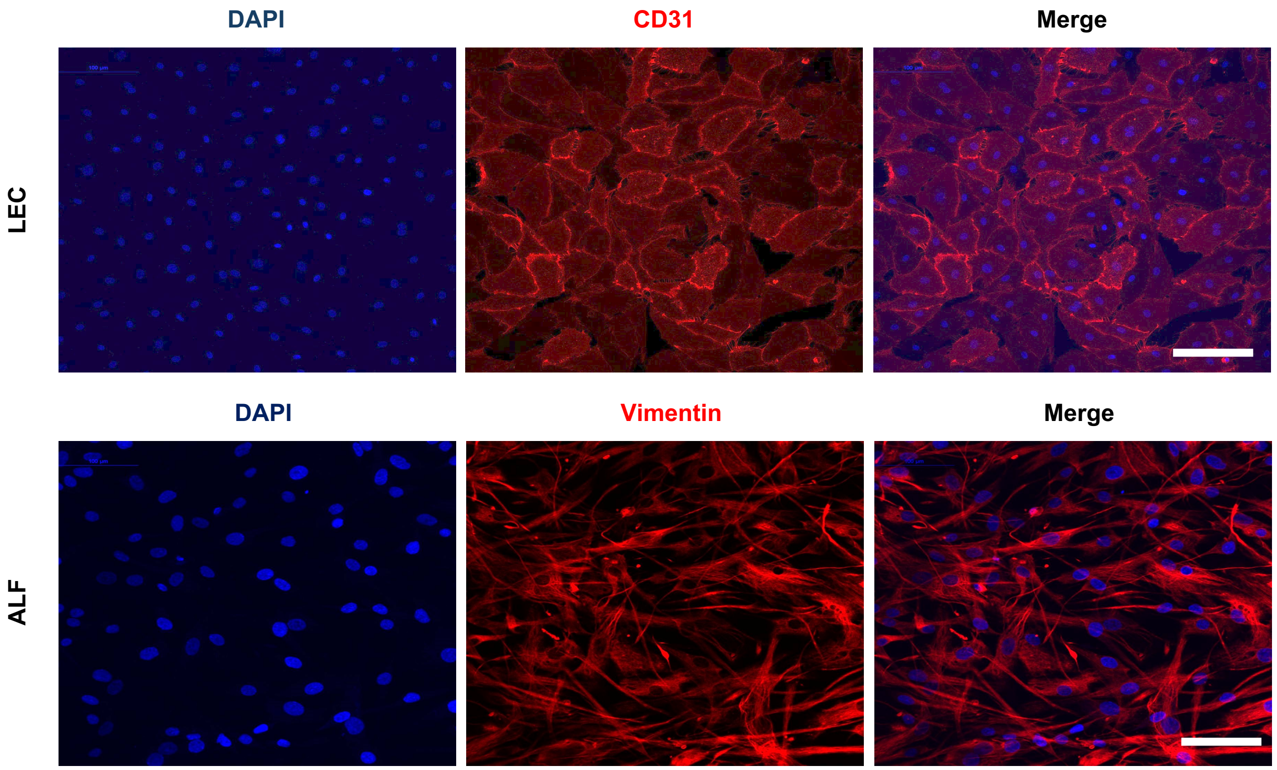


**Supplemental Fig.1: Characterization of Lung endothelial cells (LEC) and Adult lung fibroblasts (ALF).** Representative immunofluorescence images show the expression of CD31 and vimentin. Scale bar = 100 μm.

**Supplemental Fig. 2**

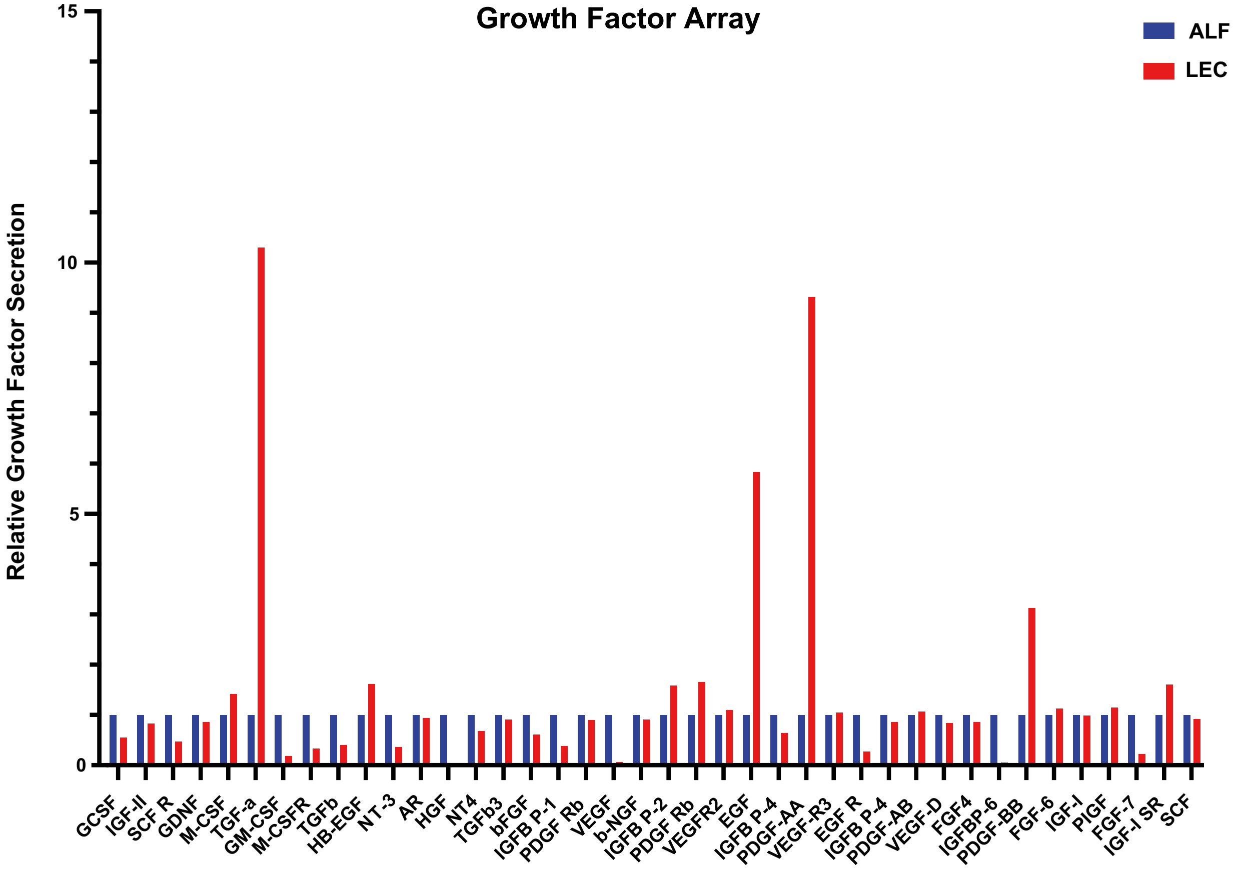


**Supplemental Fig. 2: Comparison of secreted growth factors from human lung endothelial cells (LECs) and adult lung fibroblasts (ALF).** Graphs show the relative expression of proteins detected in the growth factor array from conditioned media collected from LECs and ALF.

**Supplemental Fig. 3**

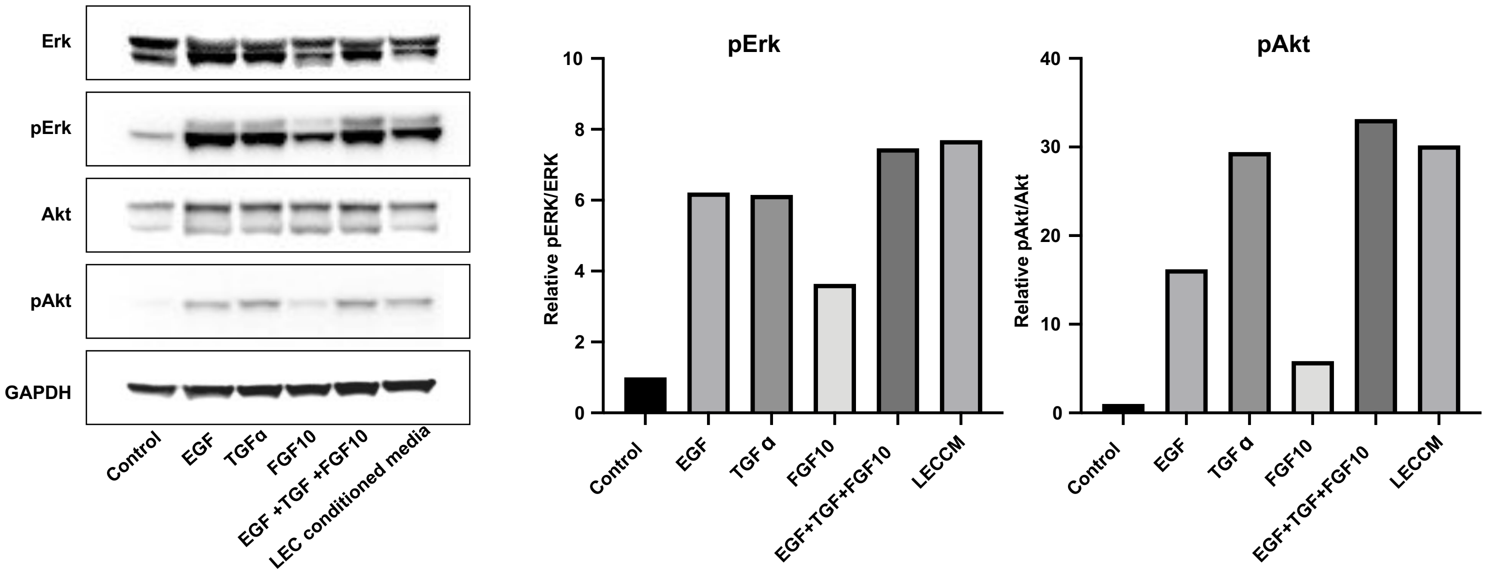


**Supplemental Fig. 3: EGF, TGFα, FGF10, and LEC conditioned media promote phosphorylation of ERK and AKT.** Western blot analysis of lysates from AT2 cells exposed for 15 min to each growth factor or lung endothelial cell conditioned media (LECCM). Graphs show relative levels of phosphorylation.

**Supplemental Fig. 4**

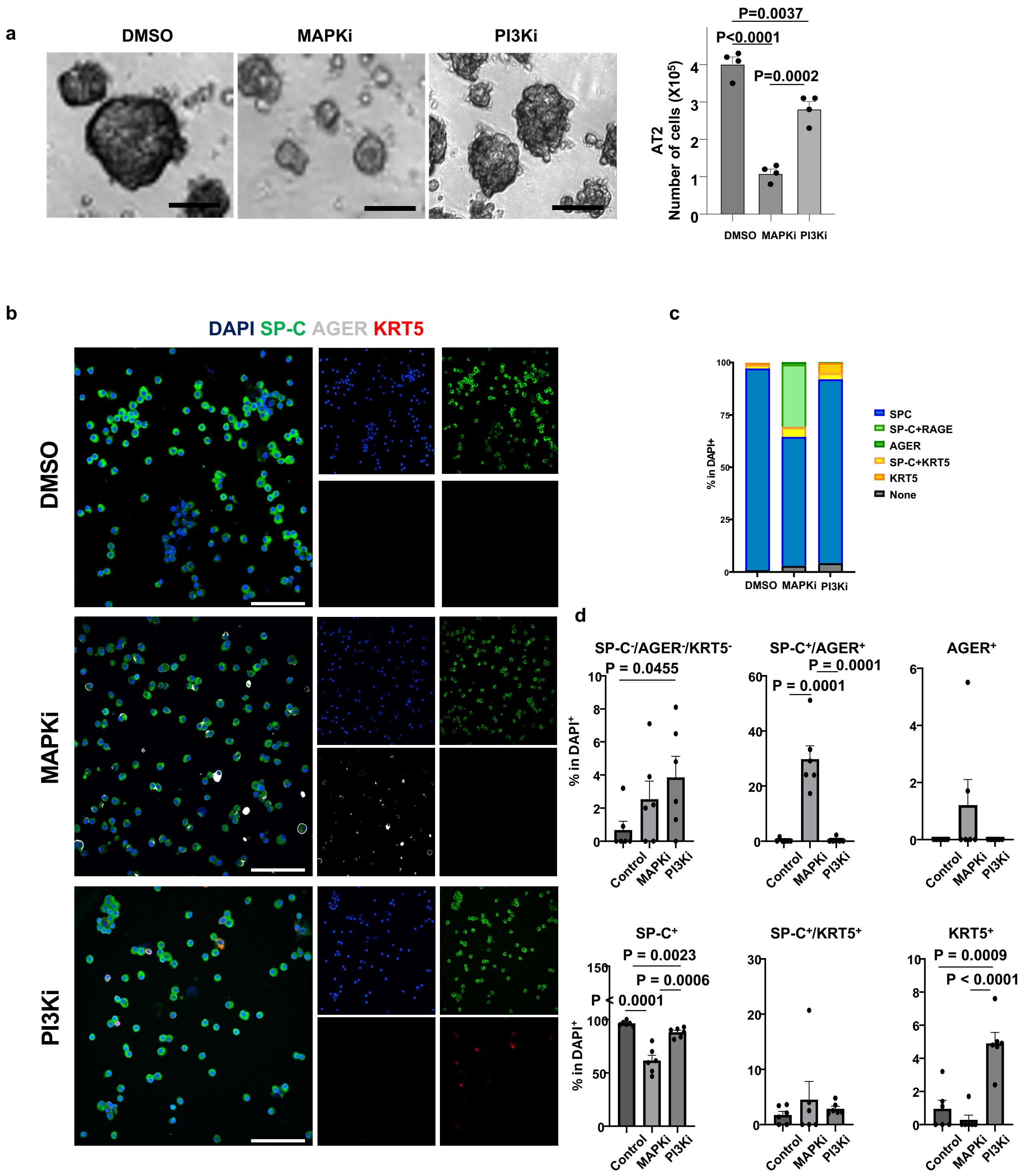


**Supplemental Fig. 4: Inhibition of MAPK and PI3K suppresses AT2 cell self-renewal and induces differentiation.** a. Phase images of AT2 cells with indicated treatment. Graph shows the number of viable cells after culture. b. Immunofluorescence staining of SP-C, AGER, and KRT5 in recovered cells. Scale bar = 100 µm. c–d. Graphs show the percentage of cells expressing each marker protein (n = 6). Statistical significance was determined by Student's t test, and data are represented as mean ± SEM.
